# Substrate sequence selectivity of APOBEC3A implicates intra-DNA interactions

**DOI:** 10.1101/176297

**Authors:** Tania V. Silvas, Shurong Hou, Wazo Myint, Mohan Somasundaran, Brian A. Kelch, Hiroshi Matsuo, Nese Kurt Yilmaz, Celia A. Schiffer

## Abstract

The APOBEC3 (A3) family of human cytidine deaminases is renowned for providing a first line of defense against many exogenous and endogenous retroviruses. However, the ability of these proteins to deaminate deoxycytidines in ssDNA makes A3s a double-edged sword. When overexpressed, A3s can mutate endogenous genomic DNA resulting in a variety of cancers. Although the sequence context for mutating DNA varies among A3s, the mechanism for substrate sequence specificity is not well understood. To characterize substrate specificity of A3A, a systematic approach was used to quantify the affinity for substrate as a function of sequence context, length, substrate secondary structure, and pH. We identified the A3A ssDNA binding motif as (T/C)TC(A/G), and found that A3A binds RNA in a sequence specific manner. Furthermore, A3A bound tighter to its substrate binding motif when in a loop compared to linear oligonucleotide. Our results suggest that the A3A affinity and preference for substrate is modulated by the structure of DNA, and not just its chemical identity. Analysis of previously published co-crystal structures of A3A bound to ssDNA in light of the above findings directed the proposal of a new model for the molecular mechanism underlying A3A sequence preference. On a broader scale, the results of this work not only provide key insights into the mechanism of A3’s beneficial roles in the cell, especially in viral restriction, but also into A3’s deleterious activity such as in the development of cancer.

## INTRODUCTION

The APOBEC3 (short for “apolipoprotein B mRNA editing enzyme, catalytic polypeptide-like”) family of human cytidine deaminases provides a first line of defense against many exogenous and endogenous retroviruses such as HIV-1 and retro-element LINE-1 (1-6). APOBEC3 (A3) proteins restrict replication of retroviruses inducing hypermutations in the viral genome (7) by deaminating the deoxycytidines in ssDNA into uridines during reverse transcription. This results in G to A hypermutations, as adenosines are transcribed across from uridines during second strand DNA synthesis. While all A3 enzymes deaminate deoxycytidines in ssDNA, they have differential substrate specificities that are context dependent, resulting in altered frequencies of mutation for the deoxycytidines. Some A3s deaminate the second deoxycytidine in a sequence containing <u>C</u>C while others deaminate deoxycytidine in a T<u>C</u> context (8-10). However, not every cognate dinucleotide motif (C<u>C</u> or T<u>C</u>) in the ssDNA of the HIV genome is deaminated (11). Nevertheless, hypermutation in a viral genome results in defective proteins and proviruses, thus decreasing the probability of further viral replication (12).

Beyond restricting viral replication, the ability of A3s to deaminate deoxycytidines in ssDNA have made A3s a double-edged sword. When overexpressed, A3s can mutate the host genome resulting in a variety of cancers. The identities and patterns of the mutations observed in cancer genomes can define the source of these mutations. Recently, the search for the deaminase(s) responsible for kataegic mutations found in breast cancer was narrowed down to APOBEC3B, through the comparison of all known APOBEC mutational signatures and eliminating APOBEC3G and other deaminases from potential mutational contributors (9, 13). Soon after, APOBEC3B was found to be correlated with a variety of other cancers such as ovarian, cervical, bladder lung, head and neck; signature sequence analysis was also a contributing factor that led to these conclusions (14, 15). Most recently APOBEC3H, which has a different sequence preference than APOBEC3B, has been identified to also play a role in breast and lung cancer (16). Thus, defining A3 sequence specificity can be helpful in identifying A3’s role in viral restriction and in cancer.

A3 signature sequences proposed for deaminating deoxycytidines range between di-nucleotide to quad-nucleotide motifs (8-11, 16-21). Although A3s are known to have varied sequence preference, quantitative and systematic studies of sequence specificity are incomplete. Recently, crystal structures of APOBEC3A (A3A) and APOBEC3B-CTD (an active site A3A chimera) with ssDNA have been solved (20, 22). However, despite these breakthrough structures the molecular mechanism underlying substrate sequence specificity flanking the TC dinucleotide sequence remains unclear.

A3A is a single-domain enzyme with the highest catalytic activity among human APOBEC3 proteins (23) and a known restriction factor for the retroelement LINE-1 and HPV (24, 25). A3A can also contribute to carcinogenesis with increased expression or defective regulation (26). A3A is the only A3 where both the intact apo and substrate bound structures have been determined (19, 20, 22, 27, 28). Initial substrate specificity studies have shown a preference for DNA over RNA, suggested by NMR chemical shift perturbation (19). Since A3A is the best biochemically characterized A3 human cytidine deaminase and thus a critical benchmark within the family, we chose A3A to elucidate the extended characteristics of ssDNA specificity.

To determine the substrate specificity of A3A, we systematically quantified the affinity of A3A for nucleic acid substrates as a function of substrate sequence, length, secondary structure, and solution pH. We identified the A3A preferred ssDNA binding motif, (T/C)TC(A/G), and found that A3A can bind RNA in a sequence specific manner. Surprisingly, A3A’s signature sequence was necessary but not sufficient to account for A3A’s high affinity for ssDNA. Significantly, A3A bound more tightly to the motif in longer oligonucleotides, and in the context of a hairpin loop. Using recently published structures of A3As complexed with ssDNA from our lab and others, we propose a structural model for the molecular mechanism for this enhanced affinity where inter-DNA interactions contribute to A3A recognition of the cognate sequence. This model provides insights into how the nucleotides flanking the canonical TC sequence may contribute to substrate sequence preference of A3A.

## MATERIAL AND METHODS

### Cloning of APOBEC3A E72A overexpression construct

The pColdII His-6-SUMO-A3A(E72A) was constructed by first cloning the SUMO gene from pOPINS His-6-SUMO into pColdII His-6 vector (Takara Biosciences) using NdeI and KpnI restriction sites. Human APOBEC3A coding sequence from pColdIII GST-A3A(E72A, C171A) was then cloned into the pColdII His-6-SUMO vector with KpnI and HindIII. The C171A mutation in the A3A construct was reverted to wild type residue by site directed mutagenesis resulting in the pColdII His-6-SUMO-APOBEC3A(E72A) catalytically inactive over-expression construct used for all experiments in this study.

### Expression and purification of APOBEC3A E72A

Escherichia coli BL21 DE3 Star (Stratagene) cells were transformed with the pColdII His-6-SUMO-APOBEC3A(E72A) vector described above. The E72A mutation was chosen to render the protein inactive. Expression occurred at 16 °C for 22 hours in lysogeny broth medium containing 0.5 mM IPTG and 100 μg/mL ampicillin. Cells were pelleted, re-suspended in purification buffer (50 mM Tris-HCl [pH 7.4], 300 mM NaCl, 1 mM DTT) and lysed with a cell disruptor. Cellular debris was separated by centrifugation (45,000 g, 30 min, 4C). The fusion protein was separated using HisPur Ni-NTA resin (Thermo Scientific). The His6-SUMO tag was removed by means of a Ulp1 protease digest overnight at 4 °C. Untagged A3A(E72A) was separated from tag and Ulp1 protease using HisPur Ni-NTA resin. Size-exclusion chromatography using a HiLoad 16/60 Superdex 75 column (GE Healthcare) was used as a final purification step. Purified recombinant A3A was determined to be free of nucleic acid prior to binding experiments by checking OD 260/280 ratios, which was at 0.54

### Oligo source and preparation

Labeled and unlabeled oligonucleotides used in this assay were obtained through Integrated DNA Technologies (IDT). Labeled oligonucleotides used in the fluorescence anisotropy based binding assay contain a 50-TAMRA flourophore at their 5’ end and were re-suspended in ultra-pure water at a concentration of 20 μM. Unlabeled oligonucleotides used for the competition assays were resuspended in ultra-pure water to a concentration of 4 mM.

### Fluorescence anisotropy based DNA binding assay

Fluorescence anisotropy based DNA binding assay was performed as described (28) with minor alterations. A fixed concentration of 10 nM 50-TAMRA-labeled oligonucleotides was added to A3A-E72A in 50 mM MES buffer (pH 6.0), 100 mM NaCl, 0.5 mM TCEP in a total reaction volume of 150 mL per well in nonbinding 96-well plates (Greiner). For the fluorescence anisotropy based DNA binding assay with APOBEC3B-CTD E255A was performed in 50 mM Tris buffer (pH 7.4), 100 mM NaCl, 0.5 mM TCEP. The concentration of APOBEC3 was varied in triplicate wells. Plates were incubated for overnight at room temperature.

For the pH dependence experiments the buffer reagent used for testing was pH 4.0–5.0 sodium acetate, pH 5.5-6.5 MES, pH 7.0-8.0 HEPES, pH 8.5-9.0 TRIS. Assay was performed as described above. For the competition assays, a fixed concentration of 300 nM A3A(E72A) was used and unlabeled oligonucleotide of varied concentration was added from 0–6.1uM. A3A(E72A) was pre-incubated with unlabeled oligonucleotide for an hour in assay buffer, then labeled DNA was added and incubated overnight at room temperature.

For all experiments, fluorescence anisotropy was measured using an EnVision plate reader (PerkinElmer), exciting at 531 nm and detecting polarized emission at 579 nm wavelength. For analyzing data and determining Kd values, Prism (GraphPad) was used for least-square fitting of the measured fluorescence anisotropy values (Y) at different protein concentrations (X) with a single-site binding curve with Hill slope, a nonspecific linear term, and a constant background using the equation Y=(Bmax 3 X^h)/ (Kd^h + X^h) +NS3X + Background, where Kd is the equilibrium dissociation constant, h is the Hill coefficient, and Bmax is the extrapolated maximum anisotropy at complete binding.

### ^1^H NMR based A3 deaminase activity assay

Deaminase activity was determined for A3 proteins by assaying active enzyme against linear DNA substrates and measuring the product formation using ^1^H NMR. Active A3A and A3B proteins with concentrations ranging from 40-400 nM were assayed against linear DNA substrates (200 μM) in buffer with 25 mM Tris pH 7.3, 100 mM NaCl, 1 mM DTT, 0.002% TWEEN20, 5% D_2_O at 298K. Experiments were performed in a Bruker Avance III NMR spectrometer operating at a ^1^H Larmor frequency of 600 MHz and equipped with a cryogenic probe. Activity was determined from the initial rate of product formation via optimization to linear equation using MATLAB software package (Mathworks Inc., Natick, MA). Product concentration was estimated from peak integrals with Topspin 3.5 software (Bruker Biospin Corporation, Billerica, MA) using an external standard. Rate errors were estimated by Monte Carlo simulation using 100 synthetic data sets and taking the residuals of the initial fit to the experimental data as the concentration error.

### Molecular Modeling

The crystal structures of A3A bound to ssDNA (PDB ID: 5KEG and 5SWW) were used for molecular modeling (20, 22). The DNA sequence was first mutated using Coot (29). The complex structure was then prepared and minimized by ProteinPrep Wizard in Maestro (Schrödinger) at pH6.0 with other settings as default.

## RESULTS

### A3A binding to ssDNA is context dependent

To interrogate the substrate sequence preference of A3A, we systematically quantified the changes in binding affinity of catalytically inactive A3A bearing the mutation E72A to a library of labeled ssDNA sequences using a fluorescence anisotropy-based DNA binding assay (28). First, to ensure that the affinity for substrate was due entirely to the sequence of interest and not due to nonspecific binding or undesired secondary structure effects, an appropriate control background sequence was identified. The dissociation constants (K_d_’s) for homo-12-mer ssDNA sequences, Poly A, Poly T, Poly C, were determined (**Figure 1A)**. Poly G was not tested due its propensity to form secondary structure elements. Poly T (750 ± 44 nM), which had previously been used in background sequences (28), bound to A3A with 2-fold higher affinity than Poly C (1,600 ± 117 nM). A3A had the lowest affinity for Poly A with a K_d_ of >11,00 nM (**Table 1**). Thus without a greater context for A3A to target, Poly C was only weakly bound. For all subsequent assays, Poly A was used, as there is no detectible binding affinity of A3A to Poly A, to provide the minimal background possible.

**Figure 1.**
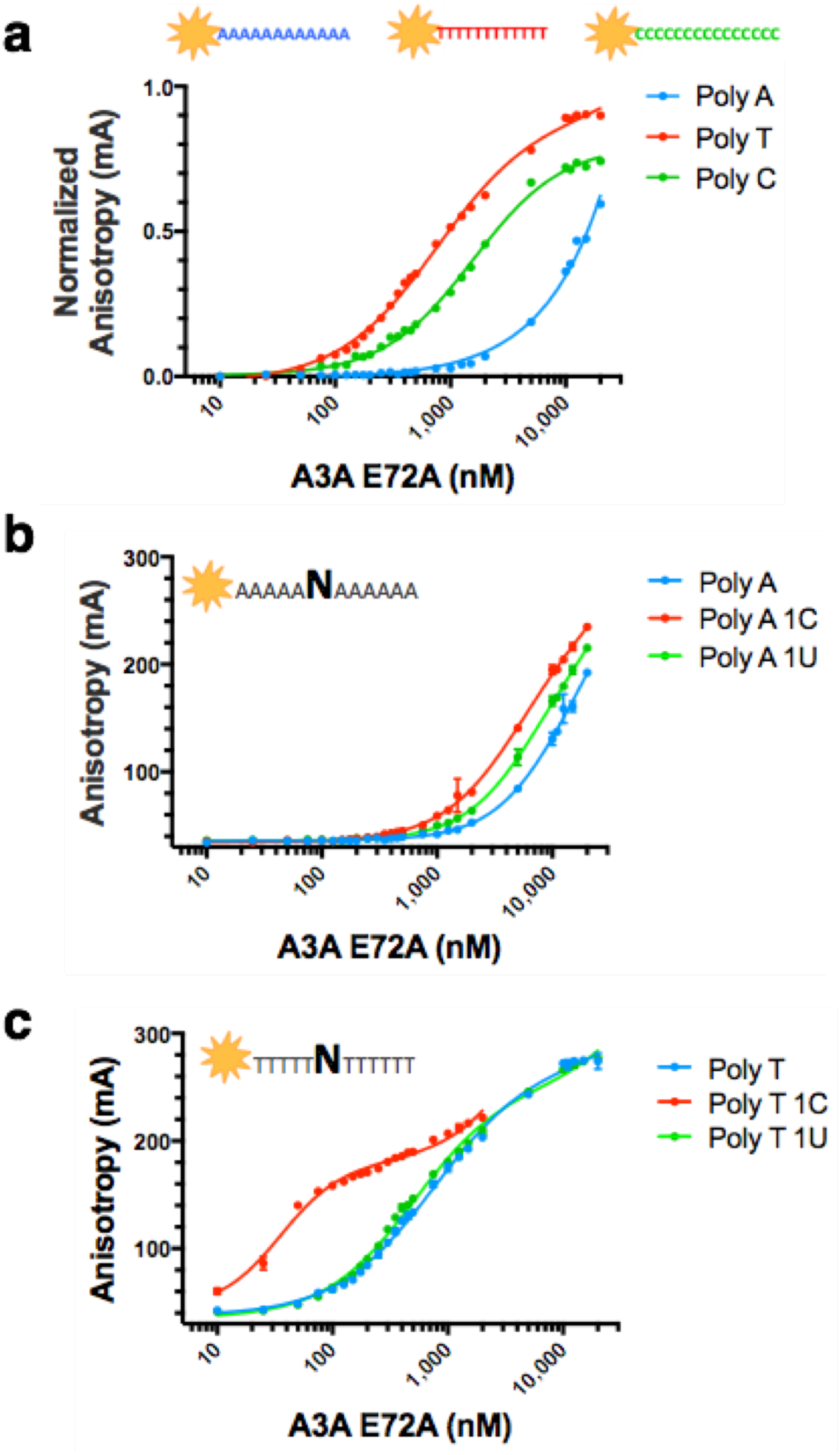
A3A specificity to ssDNA background and substrate. Fluorescence anisotropy of TAMRA-labeled ssDNA sequences binding to A3A(E72A). **a**) Binding of A3A to poly nucleotide (12 mers): Poly A (blue), Poly T (red) and Poly C (green), **b**) Binding to Poly A (blue), 5A-C-6A (red), 5A-U-6A (green), **c)** Binding to Poly T (blue), 5T-C-6T (red), 5T-U-6T (green).

**Table 1.** A3A affinity for ssDNA sequences used in this analysis.

| **DNA sequence** | **KD (nM)** |
| --- | --- |
| **Poly C (12C)** | 1,568 ± 117 |
| **Poly T (12T)** | 748 ± 44 |
| **Poly T-C (5T-1C-6T)** | 35 ± 2 |
| **Poly T-U (5T-1U-6T)** | 499 ± 23 |
| **Poly A (12A)** | >11,000 |
| **Poly A-C (5A-1C-6A)** | >5,000 |
| **Poly A-U (5A-1U-6A)** | >6,500 |
| **Poly A-TC (4A-TC-6A)** | 143 ± 4 |
| **Poly A-CC (4A-CC-6A)** | 250 ± 14 |
| **Poly A-GC (4A-GC-6A)** | >6,500 |
| **Poly A-TU (4A-TU-6A)** | >5,000 |
| **Poly A-ATCA (3A-ATCA-5A)** | 145 ± 2 |
| **Poly A-ATCT (3A-ATCT-5A)** | 209 ± 5 |
| **Poly A-ATCC (3A-ATCC-5A)** | 163 ± 3 |
| **Poly A-ATCG (3A-ATCG-5A)** | 154 ± 2 |
| **Poly A-TTCA (3A-TTCA-5A)** | 90 ± 1 |
| **Poly A-TTCT (3A-TTCT-5A)** | 127 ± 2 |
| **Poly A-TTCC (3A-TTCC-5A)** | 114 ± 2 |
| **Poly A-TTCG (3A-TTCG-5A)** | 92 ± 2 |
| **Poly A-CTCA (3A-CTCA-5A)** | 85 ± 1 |
| **Poly A-CTCT (3A-CTCT-5A)** | 122 ± 2 |
| **Poly A-CTCC (3A-CTCC-5A)** | 101 ± 2 |
| **Poly A-CTCG (3A-CTCG-5A)** | 86 ± 1 |
| **Poly A-GTCA (3A-GTCA-5A)** | 150 ± 3 |
| **Poly A-GTCT (3A-GTCT-5A)** | 218 ± 7 |
| **Poly A-GTCC (3A-GTCC-5A)** | 152 ± 2 |
| **Poly A-GTCG (3A-GTCG-5A)** | 150 ± 3 |

The specificity of A3A for substrate versus product was measured by binding to Poly A with a single C versus Poly A with a single U (**Figure 1B**). Surprisingly, the presence of a single deoxycytidine in a Poly A background was not sufficient for binding with appreciable affinity. The affinity of A3A for the Poly A-C (5A-1C-6A) (>5,000 nM) is similar to the affinity for Poly A-U (5A-1U-6A) (>6,500 nM) and even the background Poly A. This is in contrast to A3A’s specificity for binding a single C over U in a Poly T background, which is more than ten-fold (35 ± 2 nM and 500 ± 23 nM respectively) (**Figure 1C**), as we previously measured (28). This strong context dependence differentiating substrate C versus product U within the background of Poly A versus Poly T indicates that A3A heavily relies on the identity of the surrounding nucleotide sequence to recognize and bind substrate deoxycytidine.

### A3A affinity for ssDNA is pH dependent

A systematic measurement of A3A affinity in a broad range of pH values was performed to verify and quantify the pH dependence of A3A to substrate ssDNA (21, 26), and set a reference pH for subsequent experiments. The K_d_ of A3A for TTC in a Poly A background was determined at pH ranging from 4.0 to 9.0 in 0.5 pH increments (**Table 2**). A3A had the highest affinity for Poly A-TTC at pH 5.5 with a K_d_ of 68 ± 3 nM. The isotherms for A3A binding ssDNA at pHs below 6.0 show some secondary binding event that may be due to non-specific binding or aggregation (**Supplementary Figure 1b**). A steady decrease was also observed for the affinity of A3A for ssDNA when pH was increased above 6 (**Supplementary Figure 1a**), in agreement with decreased deamination activity at higher pH (26). We tested the deamination activity against two oligonucleotides at the physiological pH of 7.4, in comparison to A3B-CTD. A3A had robust deamination activity at pH 7.4 against both TCA and TCG containing ssDNA with reaction rates more than an order faster compared to A3B-CTD, in agreement with the higher substrate affinity of A3A (**Supplementary Figure 2**). A3A affinity also overall correlated with reported deamination activity determined using a different assay at pH 7.5 (30). Interestingly, A3A had no appreciable affinity for Poly A-TT<u>C</u> above pH 8.0. Since A3A is stable at these higher pH values, the lower affinity for ssDNA with increased pH is likely not due to aggregation but due to the protonation of His 29, as previously described (26) and reported to be responsible for coordinating ssDNA (31). Therefore, all of the subsequent binding experiments were performed at pH 6.0 to avoid any potential for secondary binding events or aggregation of the protein.

**Table 2.** A3A affinity for ssDNA Poly A-TTC in a range of pHs.

| **pH** | **KD** |
| --- | --- |
| **4.0** | Could not calculate |
| **4.5** | Could not calculate |
| **5.0** | Could not calculate |
| **5.5** | 68 ± 3 |
| **6.0** | 90 ± 1 |
| **6.5** | 146 ± 8 |
| **7.0** | 260 ± 21 |
| **7.5** | > 4,500 |
| **8.0** | > 5,000 |
| **8.5** | no detectable binding |
| **9.0** | no detectable binding |

### Substrate recognition is dependent on thymidine directly upstream of target deoxycytidine, with preference for pyrimidines over purines

To study the effect of the nucleotide identity at position -1 relative to target deoxycytidine (NC) on A3A affinity for substrate (**Figure 2A**), the K_d_ values of A3A for (4A)-T<u>C</u>**-**(6A), A<u>C</u>, C<u>C</u>, G<u>C</u> in a Poly A background were determined. A preference for T<u>C</u> (143 ± 4 nM), followed by C<u>C</u> (250 ± 14 nM) was identified. Interestingly, A<u>C</u> and G<u>C</u> had similarly very weak binding affinities for A3A (>5,000 and >6,500 nM respectively), illustrating a preference for pyrimidines (T or C) over purines (A or G) at -1 position with T as the strongest binder. Considering the preference of A3A for T<u>C</u>, and the much higher activity compared to C<u>C</u> (30), A3A’s affinity for substrate C versus product U was also determined in the context of Poly A. A3A preference for substrate (T<u>C</u>; 143 ± 4 nM) over product (TU; >5,000 nM – **Figure 2A** purple) in a Poly A background demonstrates the clear discrimination for substrate with T at -1 position of at least 35 fold. Thus, the smaller size and coordination of pyrimidines at -1 position appears to be a driving factor for recognizing substrate.

**Figure 2.**
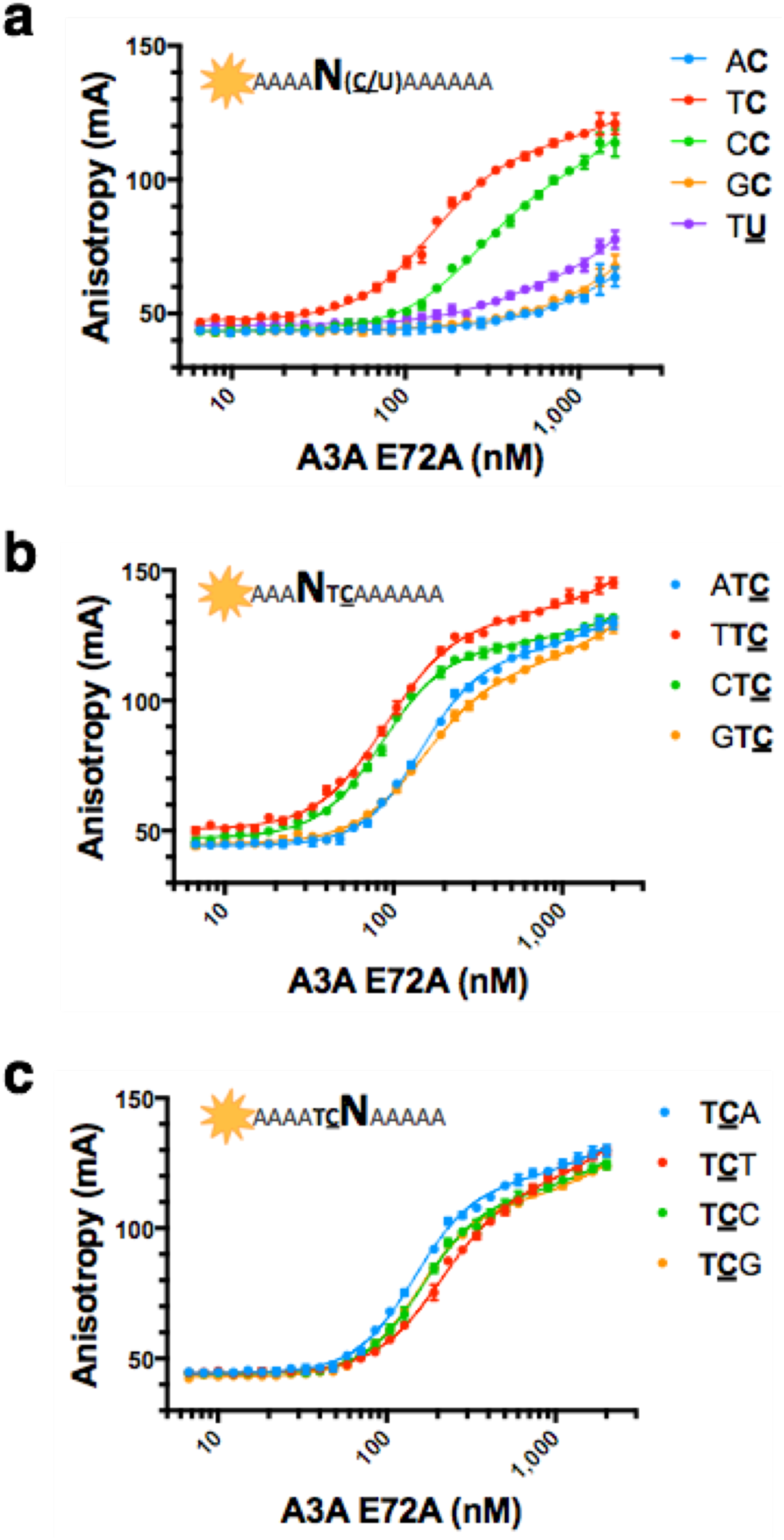
A3A specificity for nucleotides flanking substrate cytidine. Fluorescence anisotropy of TAMRA-labeled ssDNA sequences to A3A(E72A). **a**) Binding of A3A to ssDNA with changes at -1 position of substrate C and TU (purple) in a poly A background (12 mers): 4A-AC-6A (blue), 4A-TC-6A (red), 4A-CC-6A (green), and 4A-GC-6A (orange). **b**) Binding of A3A to ssDNA with changes at -2 position in a TC context in a Poly A background (12 mers): 4A-ATC-6A (blue), 4A-TTC-6A (red), 4A-CTC-6A (green), and 4A-GTC-6A (orange). **c**) Binding of A3A to ssDNA with changes at +1 position in a TC context in a Poly A background (12 mers): 4A-TCA-6A (blue), 4A-TCT-6A (red), 4A-TCC-6A (green), and 4A-TCG-6A (orange).

### A3A consensus sequence for binding cognate dinucleotide motif TC

The effects of the sequence identity around the cognate dinucleotide deamination motif (TC) on affinity of A3A for ssDNA was determined by first testing the change in affinity for all nucleotide substitutions at -2 position (3A)-NT<u>C</u>-(6A). A3A has a preference for pyrimidine over purine at -2 position (**Figure 2B**) with TT<u>C</u> and CT<u>C</u> having similar affinities (90 ± 1 nM and 85 ± 1 nM respectively) compared to that of purines AT<u>C</u> and GT<u>C</u> (145 ± 2 nM and 150 ± 3 nM respectively). While not as strong as for -1 position, there is a preference for the smaller pyrimidines at position -2. Next, the effect of +1 position on affinity of A3A to T<u>C</u> was determined. A3A did not demonstrate a strong preference for any particular nucleotide, although disfavoring T, at the +1 position (145 ± 2 nM for background versus 209 ± 5 nM).

Finally, to identify if there was any interdependency between nucleotide identity at -2 and +1 positions, the affinity of A3A for (3A)-NT<u>C</u>N-(5A) was determined (**Figure 3, Table 1**). A3A displayed preference for pyrimidines at -2 position regardless of the nucleotide at +1. A3A also disfavored T at +1 position regardless of the nucleotide identity at -2. Most interestingly, A3A preferred a pyrimidine at -2 when there was a purine at +1 position. However, the reverse was not true; purine at -2 position with pyrimidine at +1 position did not result in comparable affinities. In fact, the worst binders (AT<u>C</u>T and GT<u>C</u>T) were those that contained purines at -2 with pyrimidines at +1 position. Thus, we have identified (T/C)T<u>C</u>(A/G)as the preferred sequence for ssDNA recognition by A3A.

**Figure 3.**
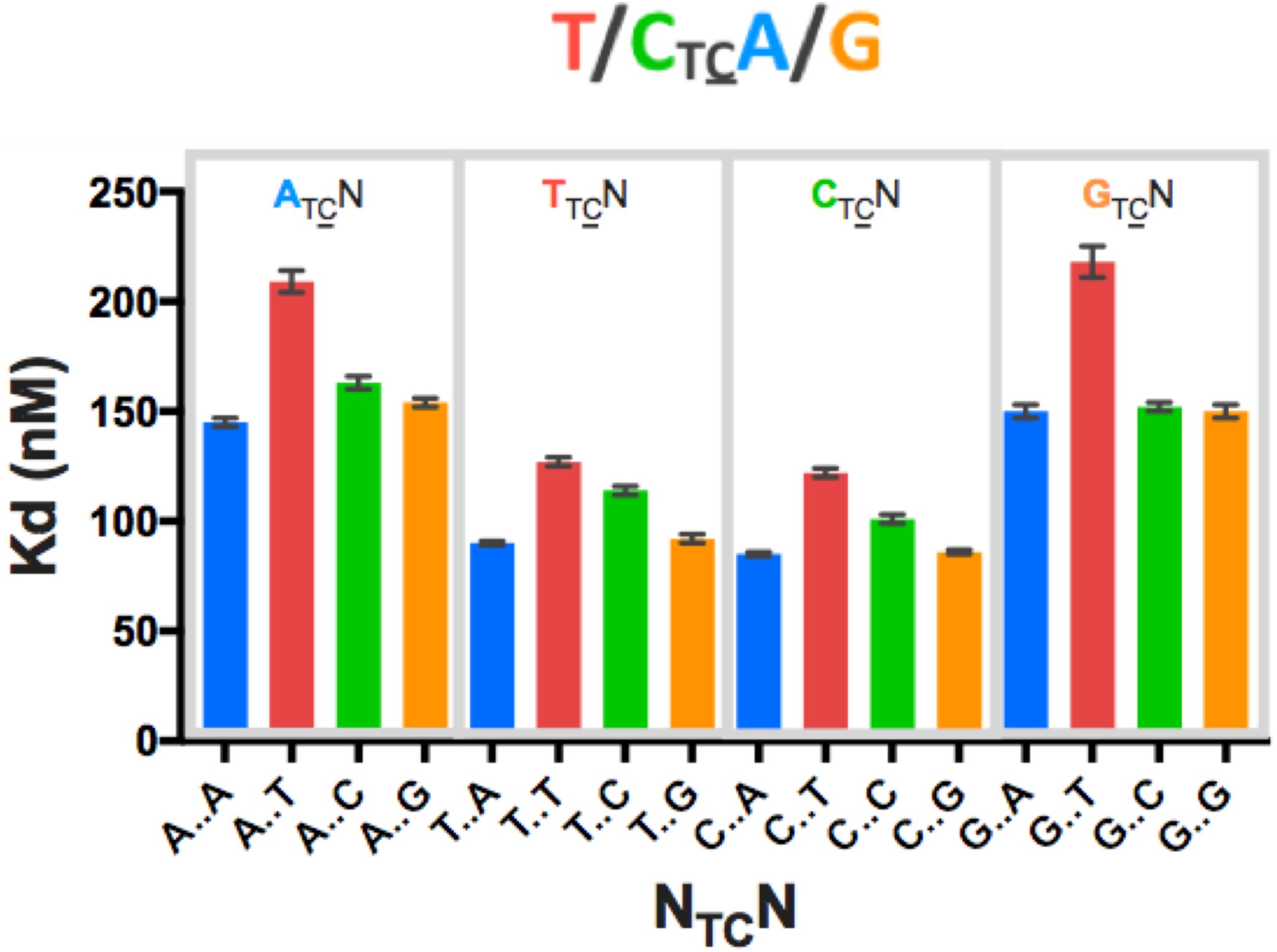
A3A specificity for poly A xTCx. Binding affinity of A3A(E72A) to TAMRA-labeled ssDNA sequences in a Poly A background. Gray boxes bin sequences by -2 nucleotide identity. Colors represent +1 nucleotide identity: A (blue), T (red), C (green), G (orange). Consensus sequence derived from these Kd values is shown above the graph.

### Structural basis for A3A specificity for binding to preferred recognition sequence

To determine the structural basis for the A3A consensus sequence (T/C)T<u>C</u>(A/G), crystal structures of A3A bound to ssDNA recently determined by our group and others (PDB ID: 5KEG and 5SWW) were analyzed. The target deoxycytidine is well coordinated and buried within the active site of A3A (**Figure 4A**) in these structures. The thymidine at position -1 has extensive contacts with loop 7 (Y130, D131 and Y132), and van der Waals contacts with loop 5 (W98) (**Figure 4B**). The Watson-Crick edge of the thymidine base faces the loop 7 residues, and makes three hydrogen bonds: one with the backbone nitrogen of Y132 and the other two, one is water mediated, are with the D131 sidechain. The D131 side chain further forms a salt bridge to the R189, which stabilizes the overall hydrogen-bonding configuration of loop 7 to the thymine base. This coordination appears critical, as residue 189 is conserved as a basic residue (Arg/Lys) in catalytically active A3 domains. At the -1 position, deoxycytidine could form altered interactions, as the N3 atom lacks the proton to hydrogen bond with D131 (**Figure 4C**). In contrast, loop 7 of A3A, in particular residues Y130 and D131, would likely preclude a larger purine base from fitting in this position, thus defining the T/C specificity at the -1 position (**Figure 4D**).

**Figure 4.**
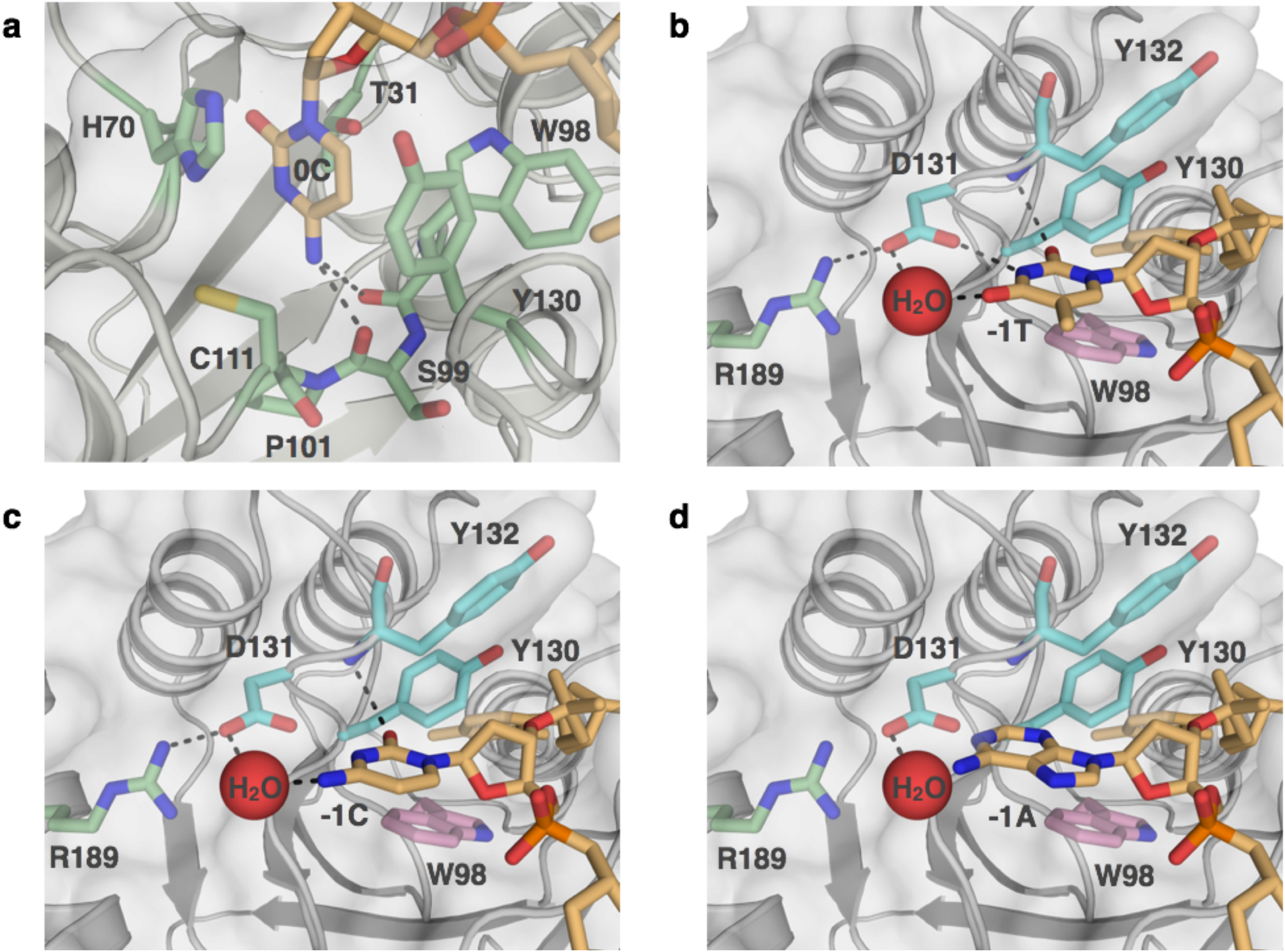
A3A recognition of substrate cytidine and pyrimidines at -1. Crystal structure of A3A(E72A/C171A) shown in surface view (gray) bound to Poly T-1C ssDNA sequence represented as sticks (PDB ID: 5KEG). **a)** substrate cytidine (orange sticks) is buried in active site of A3A. Residues interacting with cytidine are shown in green sticks. **b)** -1 nucleotide thymidine (orange sticks) surrounded by Y130, D131 and Y132 of loop 7 (light blue sticks), W98 of loop 5 (pink sticks), and R189 (green sticks). **c)** Cytidine modeled into -1 position (orange sticks). N3 atom lacks proton to hydrogen bond with D131. **c)** Adenosine modeled into -1 position (orange sticks). Other nucleotides are shown as orange sticks. Hydrogen bond and a salt bridges shown in dashes black lines. Water shown as red spheres. Nitrogen and oxygen of residues and nucleic acids are in blue and red respectively.

Although A3A has prefers (T/C)T<u>C</u>(A/G), neither of the co-crystal structures has the optimal nucleotide identity at the -2 and +1 positions. However, in both structures, the base at +1 (pyrimidine T in 5KEG and a purine G in 5SWW) stacks with the critical histidine 29 (**Figure 5A & B**) (20, 22). This type of histidine π-π stacking can occur with either a purine or a pyrimidine. However, protonated histidine prefers to stack with a purine base over pyrimidine, with thymidine stacking being the least preferred (32). Thus the specificity for purines and the disfavoring of thymidine at the +1 position relative to substrate deoxycytidine seen in our biochemical assays (**Figure 3**) can be explained by the base stacking potential with protonated histidine 29.

**Figure 5.**
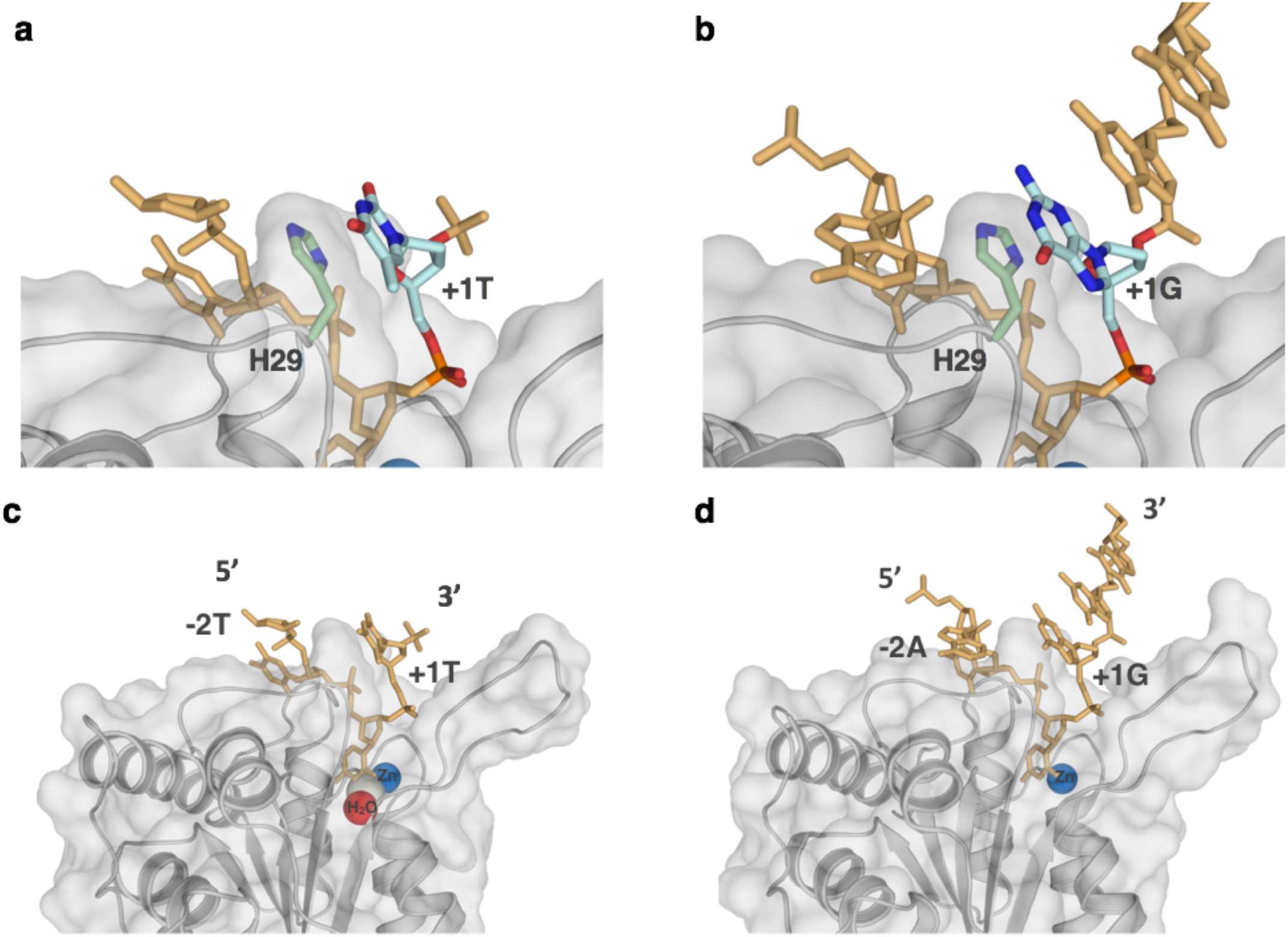
ssDNA is bent within the complex with A3A. Crystal structure of A3A shown in surface and cartoon representation (gray) bound to ssDNA displayed as orange sticks; a) +1 thymidine (light blue) is interacting with His 29 (light green sticks) through aromatic stacking (PDB ID: 5KEG) **b)** +1 guanine (light blue) also interacting with His 29 through aromatic stacking (light green sticks) (PDB ID: 5SWW). c) A3A(E72A/C171A) with TTTTTTTTCTTTTTT (PDB ID: 5KEG) **d)** A3A(E72A) with AAAAAAATCGGGAAA (PDB ID: 5SWW). Other nucleotides are shown as orange sticks, while water (red), zinc (blue), and chloride (gray) in the active site are shown as spheres. Nitrogen and oxygen of residues and nucleic acids are in blue and red respectively.

Specificity for purine at the -2 position was not evident in the available A3A–ssDNA structures. In neither structure does the -2 base make significant specific interactions, presumably as neither structure contains an optimal ssDNA sequence. For instance, even though the 5KEG structure contains a preferred pyrimidine in the -2 position, the thymidine is disordered in this complex.

A common feature between the two A3A–ssDNA complex structures is that the ssDNA forms a “U” shape in the active site (**Figure 5C & D**). This U shape of the bound polynucleotide may be conserved among deaminases, including adenosine deaminases (20, 33). In both A3A-ssDNA structures, the U shape of the ssDNA orients the -2 and +1 bases in close proximity to each other. Thus, we hypothesized that the observed sequence preference (Figure 3) for the -2 position is a result of intra-DNA interactions rather than specific interactions with the protein.

### A3A bends ssDNA to potentially allow for intra-DNA interaction between -2 and +1 nucleotides

To determine the potential for intra-DNA interactions when A3A is bound to a (T/C)T<u>C</u>(A/G) signature sequence, molecular models were developed based on the crystal structures of A3A bound to ssDNA (PDB ID: 5KEG and 5SWW). These models orient the bases of the -2 and +1 nucleotides so that they form hydrogen bonds, with the larger purine at +1 position stacking on His 29 and the smaller -2 pyrimidine coordinating the +1 base (**Figure 6**). The reversal of the nucleotides at +1 and -2 positions would not result in a fit nearly as well, which could explain the lower affinity of purine-T<u>C</u>-pyrimidine. Thus the structural model explains the preference for (T/C)T<u>C</u>(A/G) and suggests stabilizing the inter-DNA interactions may further increase the affinity..

**Figure 6.**
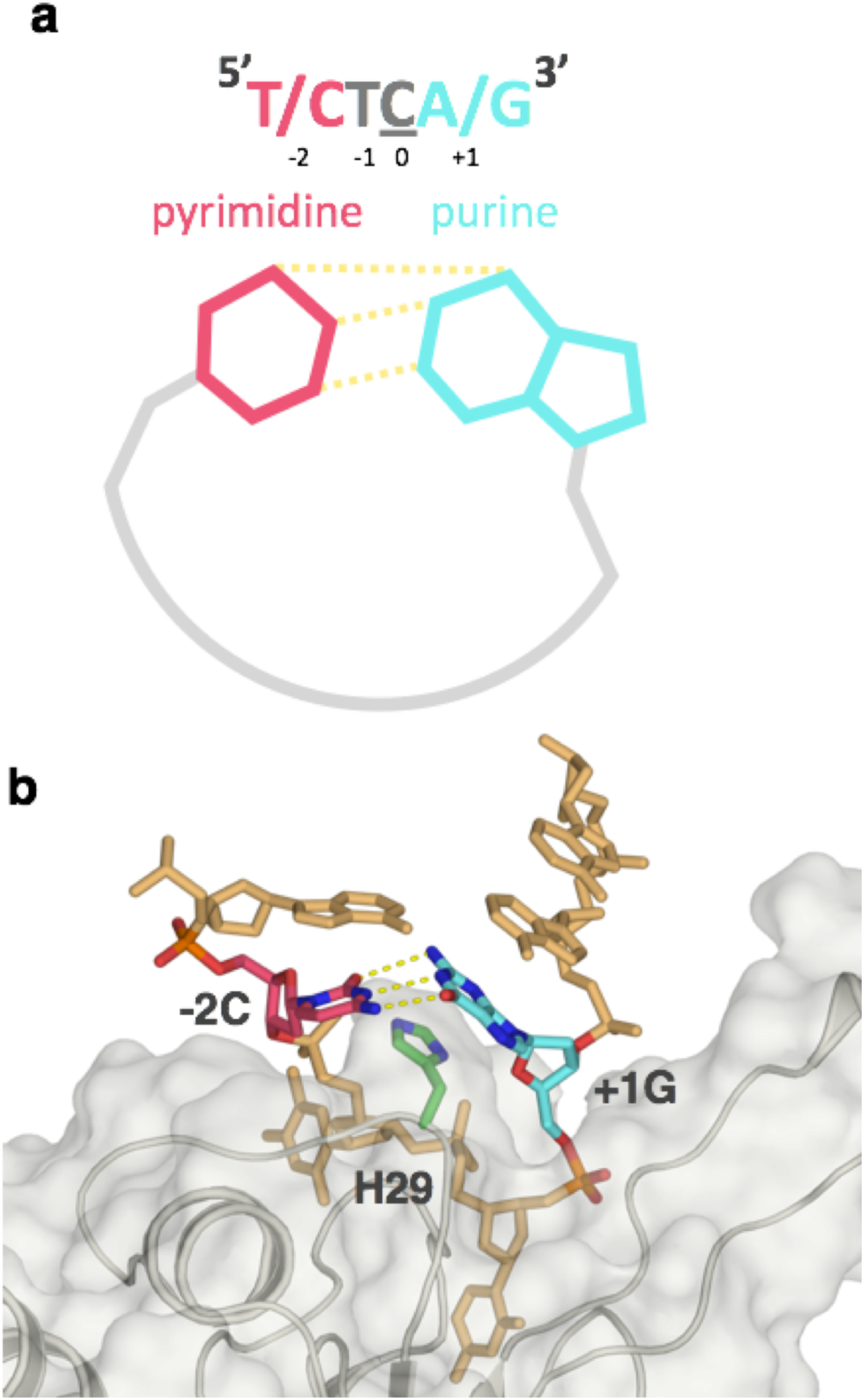
Model of inter-DNA base interactions through binding of A3A to ssDNA. **a)** A schematic of hydrogen bonding between pyrimidine (pink) at -2 and purine (light blue) at +1 position via bending of the DNA by A3A upon binding. **b)** A3A(E72A)–ssDNA complex (PDB ID: 5SWW) was used to model A3A signature sequence CTCG bound at the active site. A3A is shown as gray surface and cartoon, His29 as light green sticks, original ssDNA as orange sticks with +1G in light blue. Adenosine at -1 position was switched to cytosine (pink) with hydrogen bonds to +1G displayed as yellow dashes.

### Length of ssDNA affects affinity of A3A for substrate sequence

If the bending of the ssDNA is important for substrate recognition, dependence of binding affinity on substrate length may be expected. The length of the ssDNA that contained the recognition sequence was varied to determine if the DNA beyond the four nucleotide signature sequence contributed to the binding, in Poly A-TTC (AAA TT<u>C</u>A AAA AAA). A competition assay with different length oligonucleotides was performed to test the effect of ssDNA length on affinity for substrate (**Supplementary Figure 3**). Length was varied from 1 nucleotide flanking each end of TT<u>C</u>A (TT<u>C</u>AA and ATT<u>C</u>A) to 3 nucleotides flanking each end, increasing by one nucleotide addition on either end. Surprisingly, a single nucleotide flanking TT<u>C</u>A signature sequence was not enough to permit binding, and even three nucleotides on either side still did not bring A3A binding to original binding affinity as Poly A-TTC (AAA TT<u>C</u>A AAA AAA). Thus binding affinity is impacted beyond the recognition motif to prefer longer sequences, consistent with favoring a more complex nucleic acid structure.

### A3A prefers binding to target sequence in the loop of structured hairpins

Another implication of this model would be that pre-bent DNA could be a better substrate for A3A binding as A3A would not have to pay the entropic cost of bending the DNA. This bending of DNA could be achieved either by the inter-DNA interactions modeled in **Figure 6**, or when within a loop of a hairpin. To determine the significance of the bent U shape DNA structure in the mechanism of A3 binding, we tested A3A affinity to a target deoxycytidine in the loop region of a DNA hairpin. The hairpin was based on a previously identified hairpin sequence from succinate dehydrogenase complex iron sulfur subunit B (SDHB) RNA (34). The affinity for TTC in the loop region of hairpin DNA was higher than that in linear DNA (26 nM vs 90–127 nM respectively). As expected, A3A had a higher affinity for the DNA hairpin with loop region containing TT<u>C</u> compared to one with AAA (26 nM vs ∼676 nM respectively) (**Figure 7A**). Interestingly, the Kd value for the hairpin (26 nM) is comparable to that for a single C in a polyT background (35 nM) (28). This may imply that the polyT DNA adopts a hairpin structure in solution, as has been reported (35).

**Figure 7.**
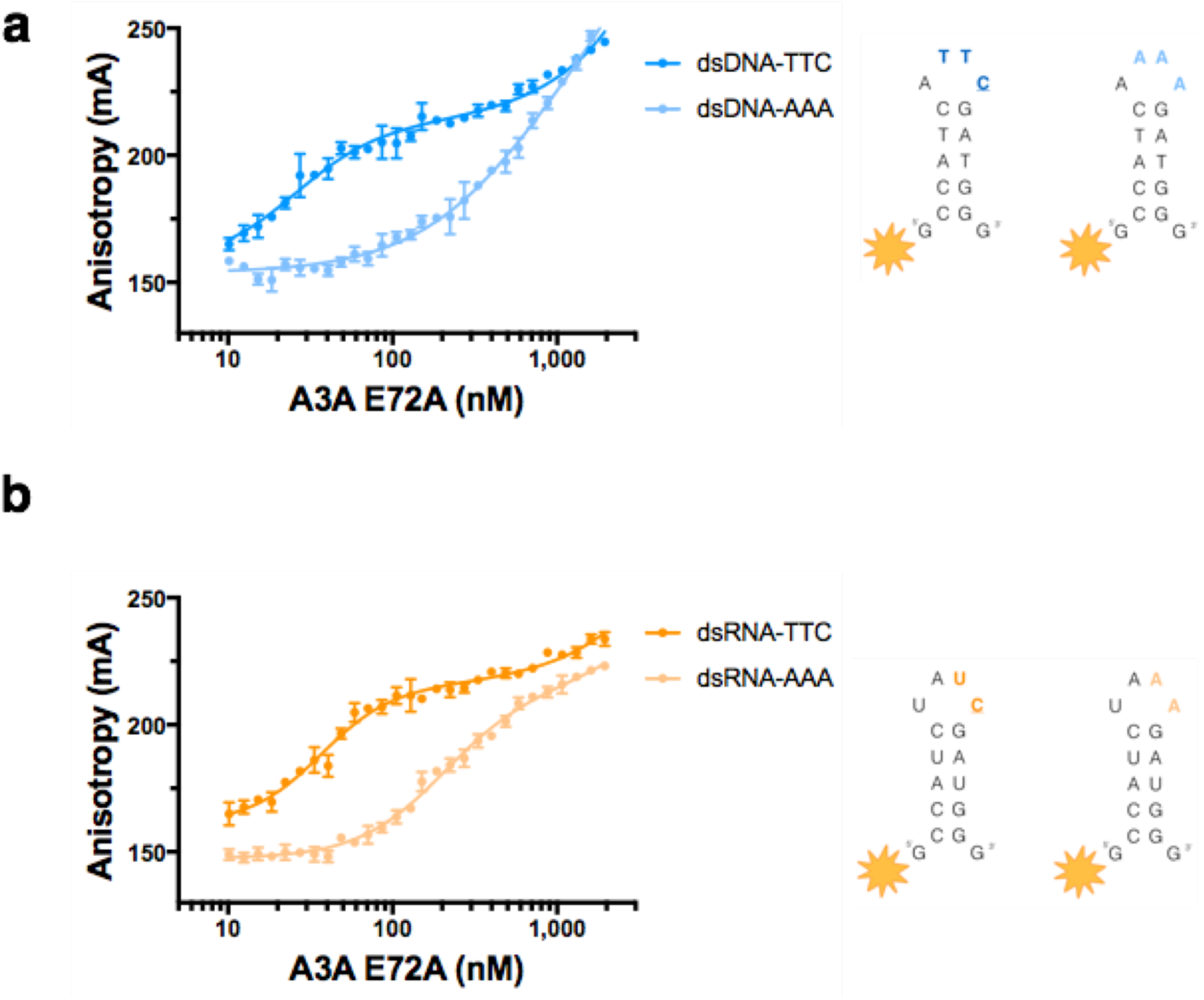
A3A specificity for substrate in loop region of stem-loop nucleic acids. Fluorescence anisotropy of TAMRA-labeled hairpin DNA and RNA to A3A(E72A). **a**) Binding of A3A to a DNA version of the hairpin SDHB RNA containing TTC (dark blue) and AAA (light blue) in the loop region. **b**) Binding of A3A to hairpin SDHB RNA (dark orange) and the same RNA sequence replacing the UC with AA in the loop region of the hairpin (light orange).

A3A affinity to a target cytidine in the loop region of an RNA hairpin was also tested. The exact SDHB hairpin RNA sequence including U<u>C</u> in the loop of this hairpin versus a modified SDHB hairpin RNA replacing the U<u>C</u> with AA was compared. A3A had specific affinity for the hairpin RNA containing U<u>C</u> compared to AA (37 nM vs 202 nM respectively) (**Figure 7B**). In contrast to what has been previously proposed (19), we found that A3A has high affinity and specificity for RNA, binding to RNA with the same interface as it does DNA. Furthermore, the affinity for AU<u>C</u> in RNA was noticeably higher than for AT<u>C</u>X in a linear sequence (37 nM vs 145–209 nM respectively). Overall, A3A has higher affinity for target sequence in the context of a pre-ordered loop region rather than linear DNA (26 nM vs 90–127 nM respectively), and specific affinity for RNA hairpins (37 nM vs 202 nM respectively).

## DISCUSSION

A3A is a single-domain enzyme with the highest catalytic activity among the human APOBEC3 proteins (23) and a known restriction factor (24, 25), which also likely contributes to carcinogenesis (26). In this study we quantified the ssDNA specificity of A3A, and identified the consensus signature sequence as (T/C)T<u>C</u>(A/G). The dinucleotide sequence preference for A3A, T<u>C</u>, which was previously found through activity assays (10, 20, 21) was confirmed and expanded to a preference for pyrimidine-T<u>C</u>-purine. Surprisingly context matters, in that the background nucleotide sequence impacts binding affinity, with essentially no binding observed for Poly A 1C **(Figure 1B**), while Poly T 1C binds with 35 ± 2 nM affinity (28). Furthermore, the length of the ssDNA in which (T/C)T<u>C</u>(A/G) is imbedded within also modulates affinity (**Supplementary Figure 3**). Structural analysis of the two A3A-ssDNA complexes containing two distinct, but suboptimal ssDNA sequences have led us to develop a model for the molecular mechanism for A3A’s specificity to ssDNA. In contrast to previous results (27), which implicate the -2 position as defining specificity, the base at this position observed in both A3A– ssDNA co-crystal structures do not make any specific interactions with the protein. Rather, the hydrogen bonding edge of the -2 base is in close proximity to corresponding edge of +1 base, prompting us to propose intra-DNA interactions as being determinants of preference. These interactions could stabilize the U-shaped DNA conformation within the A3A active site.

In this study we have also quantified A3A binding to RNA in a highly specific manner. Previous molecular reports (19) suggested that A3A bound only weakly and did not deaminate RNA. However, the potential substrate sequence was designed to lack secondary structure, which in light of our results on hairpin RNAs, may have inadvertently precluded RNA binding. Recently, A3G and A3A were implicated in deaminating RNA in proposed RNA hairpins in whole cell lysates but the specificity was not quantified (34, 36). Intriguingly, our data show that A3A binds RNA-hairpins with similar affinity as for DNA-hairpins, which suggests that RNA-editing activity from A3A might be more prevalent than previously anticipated. Future experiments will identify if A3A’s catalytic efficiency is similar for DNA and RNA-hairpins.

The comprehensive identification of A3A signature sequences and preference for loop structures will enable a more accurate evaluation of A3 activity based on sequence analysis. Previous studies used only a *single* identified A3 signature sequence to implicate A3’s role in viral restriction or cancer progression. In contrast, our study suggests a more accurate method for determining evidence of A3 activity would be to use a set of sequences. In the case of A3A, we have identified four almost equivalent substrate signature sequences, TT<u>C</u>A, TT<u>C</u>G, CT<u>C</u>A, and CT<u>C</u>G, which should be used for identifying A3A’s involvement in mutagenesis. In addition, the probability of mutagenesis should not be solely based on nucleotide sequence, but should also be weighted by the propensity of the target sequence to be within a structured loop. Secondary structure prediction software could be used to identify the consensus sequence in loop regions of structured DNA or RNA. A3A signature sequences, (T/C)T<u>C</u>(A/G), that we identified, not only accounts for the discrepancies in the A3A target sequences reported in the literature such as TT<u>C</u>A versus CT<u>C</u>G (21) (20), but also leads us to advocate a new paradigm for identifying A3A’s involvement in mutation of endogenous or exogenous DNA.

Designing inhibitors or activators for A3s has been extremely challenging. Our results implicate a need to incorporate the structural context of the target deoxycytidine in the therapeutic design. Larger “U” shaped macrocycles may serve as more appropriate starting scaffolds in designing cancer therapies targeting A3s, which would mimic the “U” shape of the bound ssDNA. Macrocycles have recently been shown to have good drug-like properties and may be a strategy to target these critical enzymes(37).

## FUNDING

This work was supported by the US National Institute of Health [R01GM118474, P01 GM091743]; and T.V.S. is supported by US National Institute of Health F31 GM11993. Funding for open access charge: US National Institute of Health.

## CONFLICT OF INTEREST

None of the material in the submitted manuscript has been published, or is under consideration for publication elsewhere. None of the authors have any commercial conflict of interest related to the content of this manuscript.

---

**Supplement Figure 1.**
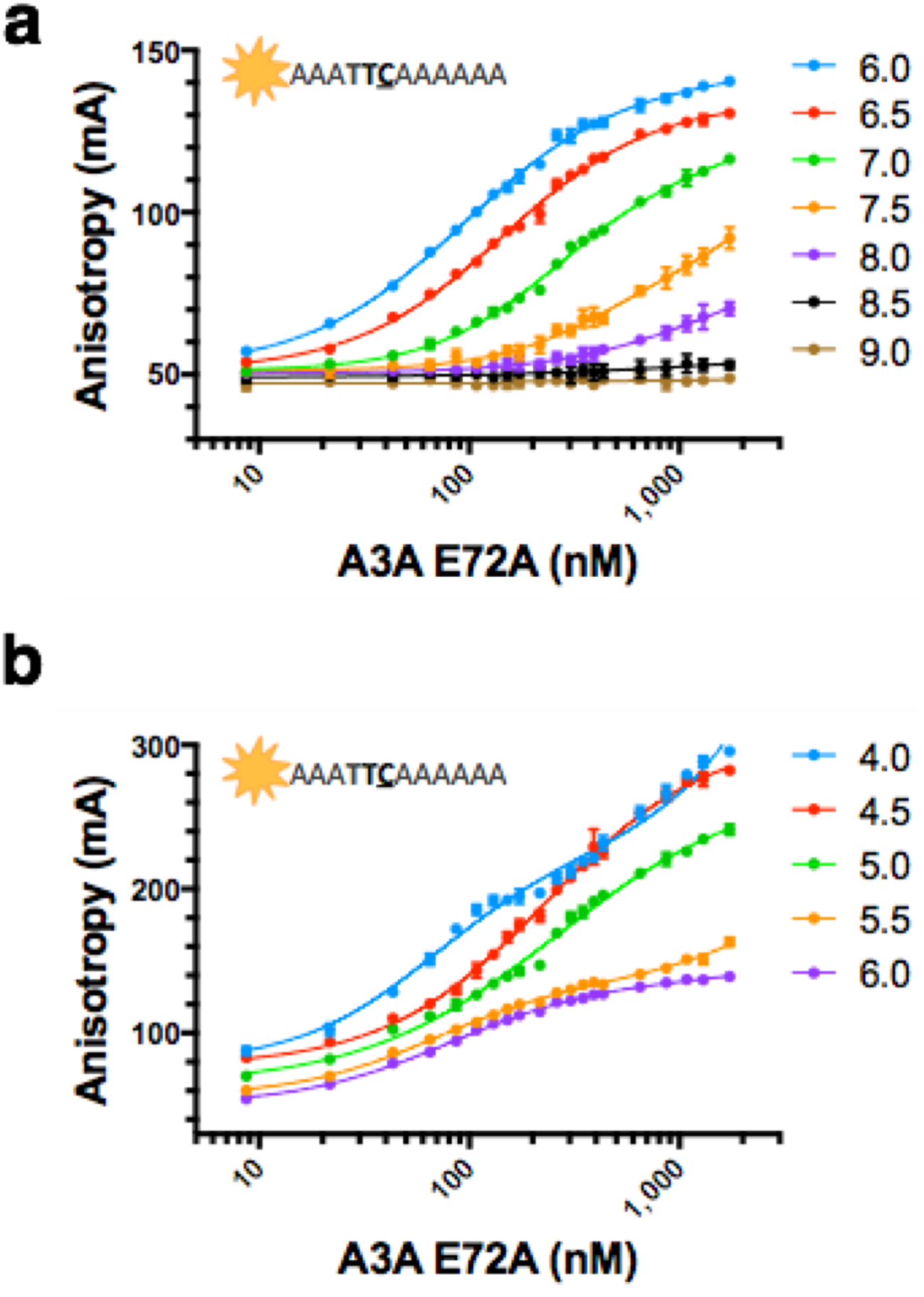
A3A affinity to ssDNA at different pHs. Fluorescence anisotropy of TAMRA-labeled ssDNA 4A-TTC-6A binding to A3A(E72A). **a)** Binding of A3A to ssDNA at pH 6.0 (blue), 6.5 (red), 7.0 (green), 7.5 (orange), 8.0 (purple), 8.5 (black), 9.0 (brown). **b)** Binding of A3A to ssDNA at pH 4.0 (blue), 4.5 (red), 5.0 (green), 5.5 (orange), and 6.0 (purple).

**Supplement Figure 2.**
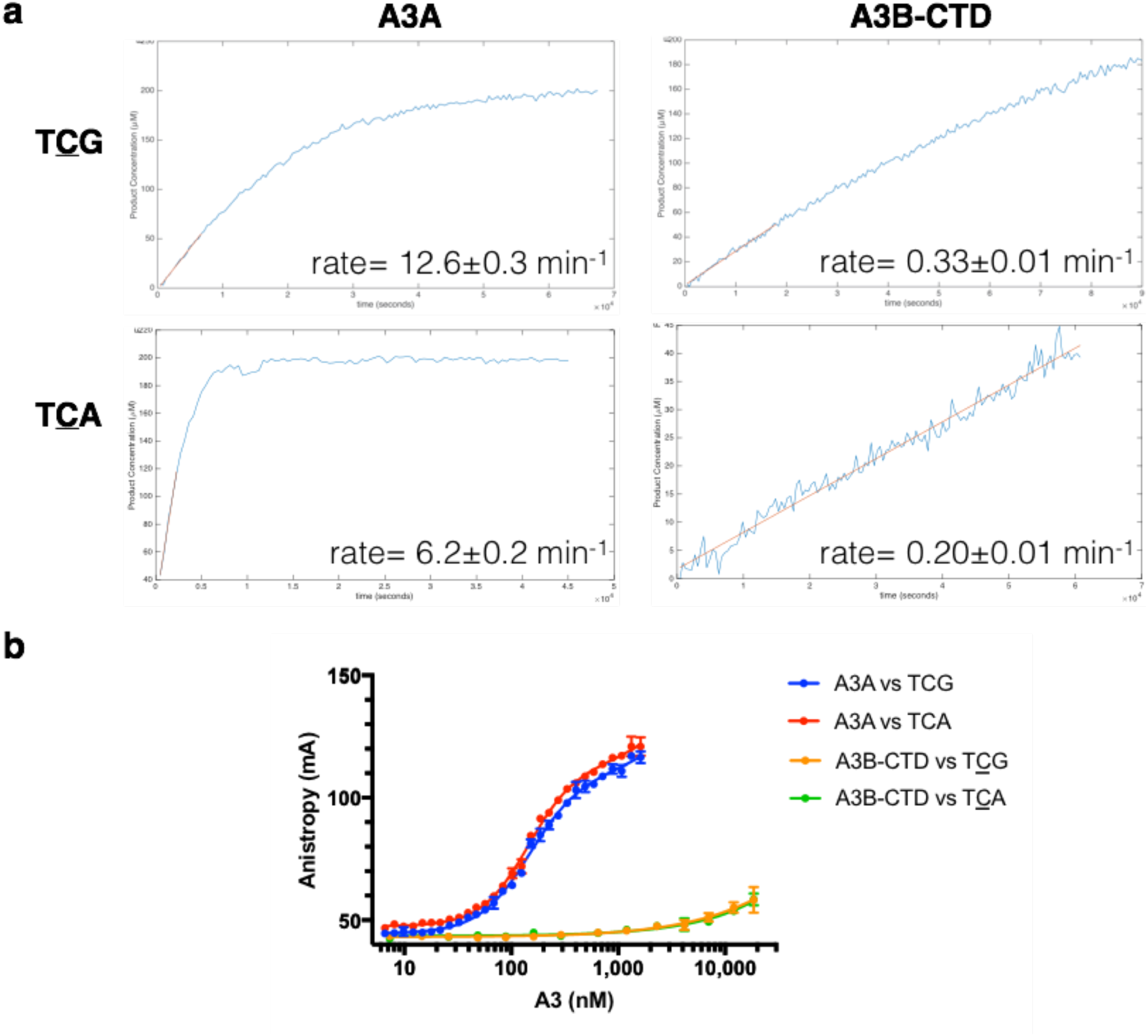
A3A and A3B-CTD deamination activity and affinity to ssDNA. **a)** Speed of deamination activity versus substrate sequence for A3A and A3B-CTD. Initial speed was determined for the conversion of 200 μM substrate of the indicated sequence at pH 7.3. The enzyme concentration for A3A versus TCG, A3A versus TCA, A3B-CTD versus TCG, and A3B-CTD versus TCA was 40, 400, 500, and 200nM respectively. Active A3A and A3B-CTD were expressed and purified as previously described for APOBEC3s (31) **b)** Fluorescence anisotropy of TAMRA-labeled ssDNA sequences to A3A(E72A) and A3B-CTD (E255A). Binding of A3A to TCG (blue) and TCA(red), and binding of A3B-CTD to TCG (orange) and TCA (green). Inactive A3B-CTD was expressed and purified as previously described for APOBEC3A (22).

**Supplement Figure 3.**
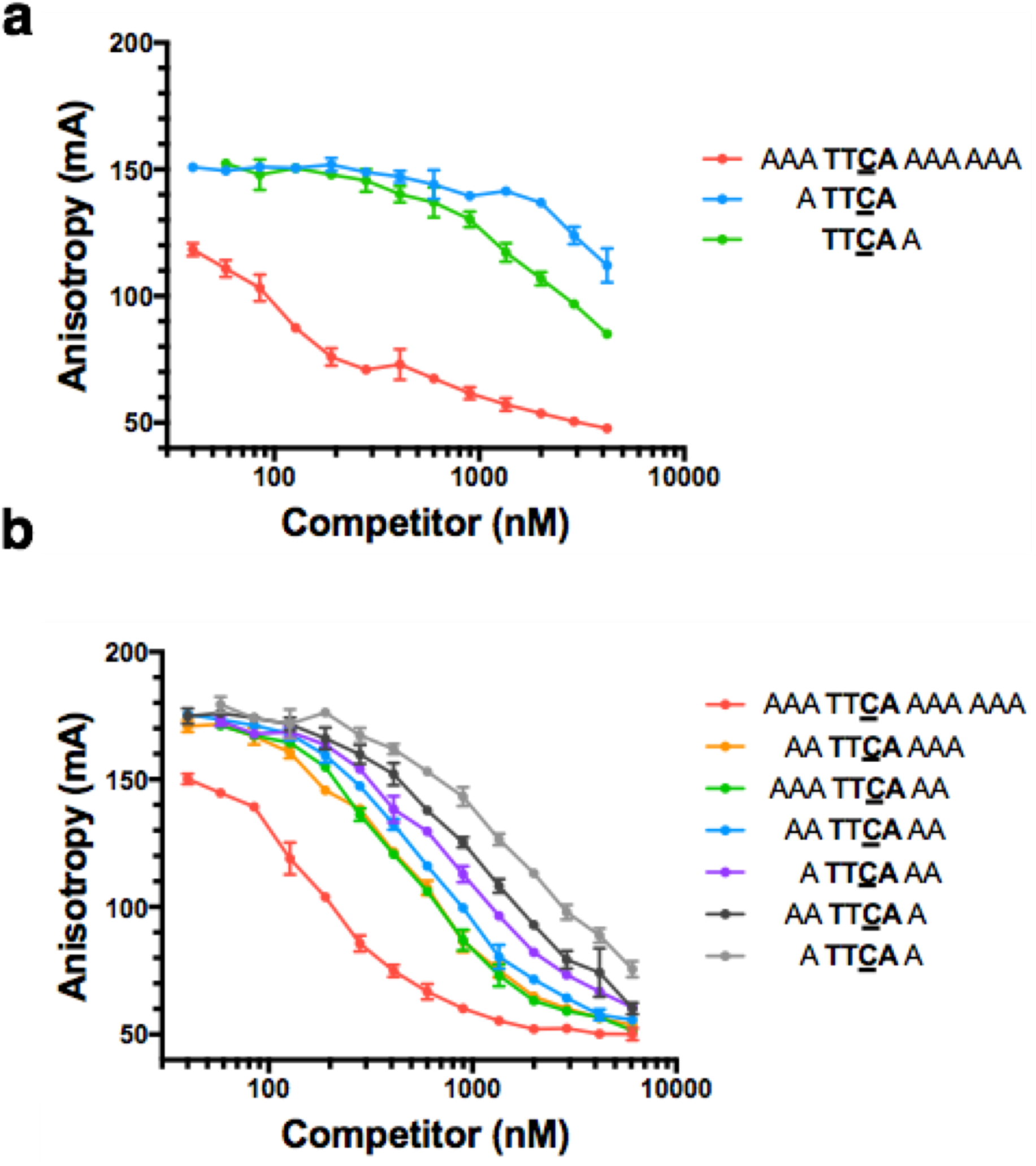
A3A affinity to ssDNA of varied lengths. Fluorescence anisotropy of TAMRA-labeled ssDNA 3A-TTCA-6A to A3A(E72A) competing with unlabeled ssDNA of different lengths. **a)** Binding of A3A to labeled ssDNA preincubated with unlabeled 3A-TTCA-6A (red), 1A-TTCA (blue), and TTCA-1A (green). **b)** Binding of A3A to labeled ssDNA preincubated with unlabeled 3A-TTCA-6A (red), 2A-TTCA-3A (blue), 3A-TTCA-2A (green), 2A-TTCA-2A (blue), 1A-TTCA-2A (purple), 2A-TTCA-1A (black), and 1A-TTCA-1A (gray).

